# Neural correlates of face perception modeled with a convolutional recurrent neural network

**DOI:** 10.1101/2023.01.02.522523

**Authors:** Jamie A. O’Reilly, Jordan Wehrman, Aaron Carey, Jennifer Bedwin, Thomas Hourn, Fawad Asadi, Paul F. Sowman

## Abstract

Event-related potential (ERP) sensitivity to faces is predominantly characterized by an N170 peak that has greater amplitude and shorter latency when elicited by human faces than images of other objects. We developed a computational model of visual ERP generation to study this phenomenon which consisted of a convolutional neural network (CNN) connected to a recurrent neural network (RNN). We used open-access data to develop the model, generated synthetic images for simulating experiments, then collected additional data to validate predictions of these simulations. For modeling, visual stimuli presented during ERP experiments were represented as sequences of images (time x pixels). These were provided as inputs to the model. The CNN transformed these inputs into sequences of vectors that were passed to the RNN. The ERP waveforms evoked by visual stimuli were provided to the RNN as labels for supervised learning. The whole model was trained end-to-end using data from the open-access dataset to reproduce ERP waveforms evoked by visual events. Cross-validation model outputs strongly correlated with open-access (r = 0.98) and validation study data (r = 0.78). Open-access and validation study data correlated similarly (r = 0.81). Some aspects of model behavior were consistent with neural recordings while others were not, suggesting promising albeit limited capacity for modeling the neurophysiology of face-sensitive ERP generation.

## Introduction

Visual perception of objects is a combined function of multiple neural systems and cognitive processes. Simply put, the visual system processes visual information received by the eyes, while the cognitive system organizes and interprets this information. These two systems work together to allow us to perceive and understand the objects and scenes we encounter in the world around us. Faces represent a special object category in human perception. Face perception has evolved in humans to be highly specialized and efficient ^1^, reflecting the central role that faces play in social interactions and communication. Humans are highly skilled at recognizing and identifying individual faces, an ability that reflects the importance of faces in our social lives and the abundance of face stimuli that we encounter in our environment. The perception of faces is the primary information source for the recognition and identification of specific individuals, is critical for interpreting expressions and emotions, and underpins the extraction of important social cues; hence, impaired visual or visual-cognitive perceptual systems can severely challenge social engagement.

In light of this, investigating the relationship between face perception and neural activity may help us to understand putative disruptions to neural systems in clinical conditions such as autism ^2^, congenital or acquired prosopagnosia ^3^, and schizophrenia ^4^, which often involve atypical face perception as measured by behavioral and neurophysiological measures.

A consistent finding from event-related potential (ERP) studies concerning face perception is sensitivity of visually evoked N170 to images of human faces ^5–8^. This ERP peak occurs earlier in response to human faces than other objects and typically has greater amplitude at right hemisphere occipitotemporal electrode sites associated with face processing specialization ^9,10^. There remains considerable debate in the literature about whether the N170 component is specific to faces or rather represents a response to highly familiar stimuli - of which faces invariably are. One argument often cited in support of the specificity view is the observation that familiar but non-face objects tend to elicit an N170 later in time than faces do. This suggests that the N170 may be more closely tied to face processing than to general familiarity with a stimulus, and supports the idea that it reflects a specific brain process devoted to face perception. However, these two explanations are not necessarily mutually exclusive, and scalp-recorded ERP signals may well reflect overlapping representations of both face specificity and more general object familiarity.

This argument requires methodological improvements to resolve. Promising, data-driven signal extraction techniques have recently become tenable through the proliferation of machine learning methods and high-performance computing. In particular, deep artificial neural networks capable of modeling complex nonlinear functions relating inputs to outputs can potentially yield valuable insights from neurophysiological data ^11,12^. For example, convolutional neural network (CNN) models have been used to investigate the computational principles underlying visual perception ^13^. These are machine learning models specialized in visual processing tasks such as object recognition. Operations performed by CNNs are considered mechanistically analogous to neurobiological implementations of vision, thus providing useful models for studying these systems ^13^. While CNNs are adept at image processing, they lack the required memory elements to handle sequential, time-varying data. Recurrent neural networks (RNNs) solve this problem with recurrent connections that provide information from previous time-steps, thereby enabling the network to model sequences. These are widely used in natural language processing, and have recently been applied to model ERP waveforms ^14–17^. Examples of RNN use in evoked response modeling include simulating ERPs to different stimuli ^14,15^, estimating changes in ERP morphology between states of consciousness ^17^, and exploring the computational processes of auditory-evoked potential generation ^14–16^.

These new analytical techniques based on machine learning algorithms can be applied to ERP data from face perception studies to help interpret the neurophysiology elicited by viewing different images. In this study we developed a computational model of human visually-evoked responses by combining the image processing capabilities of a CNN with the sequence learning functions of an RNN. We then fitted it to normative data from a classic N170 experiment. Variants of this design have been used in other contexts. For example, Shi et al. used a CNN-RNN model for action recognition in video clips ^18^, and Xu et al. used a similar architecture for epileptic seizure detection ^19^. Our approach is different insofar as we trained our model to reproduce human ERP waveforms in response to sequences of input images that were presented to subjects in a face perception study. Behavior of the model was then analyzed to investigate its correspondence with theories about ERP sensitivity to face and non-face images.

## 2. Methods

### 2.1. Data

#### 2.1.1. ERP CORE dataset

The face N170 dataset was downloaded from the ERP CORE (Compendium of Open Resources and Experiments) repository ^20^. For brevity this dataset will be abbreviated to EC. This consisted of EEG recorded from 40 healthy adult human subjects viewing a pseudo-random sequence of face (F) and car (C) images and their phase-scrambled counterparts (SF and SC) ^6^. There were 40 images in each category. All were presented twice; once in the first half and once in the second half of the sequence. Signals were band-pass filtered from 0.1 to 20 Hz then resampled to 100 Hz before re-referencing to the montage mean. Channels PO7 and PO8 were then selected for further analysis. Epochs were extracted from 0.2 s before to 0.8 s after each image presentation. Images were shown for 0.3 s in the experiment. Baseline correction was applied by subtracting the average pre-stimulus value from the whole epoch. The grand-average ERP waveform from each stimulus was produced by averaging across subjects. This resulted in 160 ERPs measured from two channels that were 101 samples in length. These provided labels for optimizing the model with supervised learning ^16^.

#### 2.1.2. Synthetic dataset

Artificial images were synthesized using generative adversarial networks (GANs) ^21^. Pre-trained StyleGAN2 models were fine-tuned with active discriminator augmentation ^22^ to generate images comparable with ERP CORE stimulus images. To prepare training data for these GANs, real images were resized to 512 px high with bi-cubic interpolation then padded to 512 px wide with picture elements matching the background gray value of 205 from the 8-bit bitmap format images. Training images were saved in 8-bit portable network graphics format. Two training sets were produced: one with 40 face images and another with 40 car images, each used to fine-tune a separate GAN. Both of the GANs had been pre-trained on the Flickr-Faces-HQ (FFHQ) dataset ^23^. We explored the same GAN architecture pre-trained on car images ^23^ to generate car images, but found that this was outperformed by the GAN pre-trained on face images. This is considered to be due to pre-training face images being higher resolution (1024 by 1024 px) than pre-training car images (512 by 384 px). Both GANs were implemented on a computer system with a graphics processing unit (Titan RTX 24 GB, Nvidia; Santa Clara, CA, USA). Batch size for each training step was 32. Fréchet Inception Distance (FID) ^24^ was computed every fourth step and training stopped if no improvement in FID was observed for 160 consecutive steps. The face-generating GAN was trained for 8900 epochs, achieving minimum FID score of 21.67; the car-generating GAN trained for 3300 epochs, achieving FID score of 50.41.

One thousand artificial face and car images were produced by the GANs after training. Truncation for StyleGAN2 was set to 0.65; the range of values this parameter can take is from 0 to 1, and it controls a trade-off between maximum fidelity (0) and diversity (1) of generated images. The resulting images were cropped and resized to match the format of original ERP CORE stimulus images. By visual inspection and FID scores these synthetic images were deemed to be comparable with but not identical to images in their respective training sets. Some synthetic face images appeared to be slightly different images of persons from the training dataset. Other synthetic face images can be described as combinations of two or more persons found in the training dataset. Synthetic car images generally looked genuine, although the interiors were relatively poorly constructed and seemed to depict a row of three front seats. Despite these and some other minor limitations, such as asymmetrical front light designs, synthetic car images were deemed adequate. Importantly, none of the artificial images were identical to any of the real images found in the training sets. These generated images were also phase-scrambled to produce a complete synthetic image dataset containing one-thousand images of each stimulus category. This data was used to investigate CNN-RNN model responses in a series of simulated experiments.

#### 2.1.3. Validation dataset

An experiment was designed to collect out-of-sample data to compare with model outputs and EC data. This design was reviewed and approved by the Macquarie University ethics committee. This dataset is abbreviated to MQ. Identical images to those used in ERP CORE ^6^ were shown to 16 healthy adult human subjects. Face (F), car (C), scrambled face (SF) and scrambled car (SC) images were presented upright and inverted (upside-down) in three different stimulus blocks counterbalanced across subjects. The first block included F, C, SF and SC conditions, equivalent to the ERP CORE sequence. The second block contained upright and inverted F and C images. The third block comprised inverted F, C, SF and SC images. Each block included 160 images played twice. Presentation^®^ (20.2, Neurobehavioral Systems Inc.; Berkeley, CA, USA) software was used to present these sequences with synchronized, event-coded time-stamps. These event codes were delayed to account for monitor refresh rate. Electroencephalography signals were recorded using a BioSemi^®^ Active Two (BioSemi Systems; Heerlen, the Netherlands) with an equivalent montage to the EC dataset. Filtering, resampling, re-referencing, channel selection, epoch segmentation, baseline correction and grand-average ERP calculation were performed as described in section 2.2.1. above.

### 2.2. Modeling

#### 2.2.1. Architecture and training

The model design can be subdivided into CNN and RNN sections illustrated in Figure 1. Input tensors were sized 101 (time-samples) by 210 (image height) by 184 (image width) by 3 (color channels). These fed into the first of ten three-dimensional convolutional layers. Each convolutional layer had 16 filters sized 1 (time-sample) by 3 (height) by 3 (width), effectively filtering over spatial dimensions of the image data while preserving its temporal structure. Filter activations were rectified and no zero padding was applied. Each pair of consecutive convolutional layers was followed by a batch normalization layer ^25^. The first four batch normalization layers were followed by three-dimensional max pooling layers ^26^ with kernel size of 1 (time-sample) by 2 (height) by 2 (width), causing tensors to compress over spatial dimensions. Outputs from the fifth batch normalization layer were reshaped to 101 (time-samples) by 240 (filter activation maps), effectively a time-sequence of vectors representing the input image sequence.

**Figure 1.**
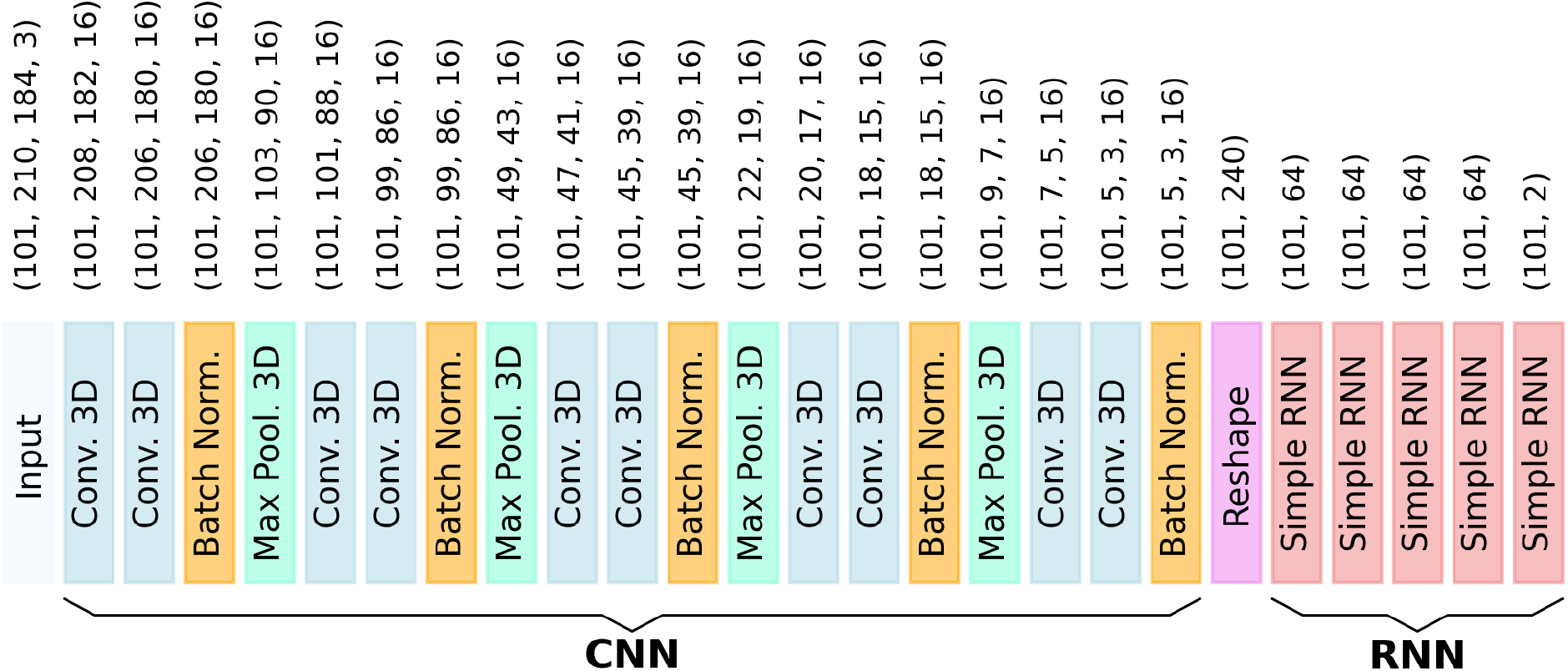
Model architecture. Convolutional neural network and recurrent neural network sections are annotated in bold. Sequentially connected layers are illustrated with colored blocks. The shape of output tensors from each layer is noted above each colored block. Convolutional layers decreased the size of spatial dimensions by two picture elements because of the filter size and absence of zero-padding. Max pooling layers decreased the spatial scale by a factor of two, rounded down for odd-numbered sizes. Temporal structure (101 time-samples) of the data is maintained throughout the network.

Sequences of vector representations from the CNN section were fed into the RNN section. The first four simple recurrent layers had 64 units with the rectified linear unit activation function applied. The fifth recurrent layer was the model output. It had two units, one for reproducing ERP data from channel PO7 and the other for PO8. The model was trained end-to-end. Input data represented visual stimuli presented to humans in the face N170 experiment, and output labels were derived from corresponding ERP waveforms. The ERP epoch spanned −0.2 s to 0.8 s around each stimulus onset time, sampled at 100 Hz. Stimulus images were shown for 0.3 s, but otherwise the screen was gray. This scenario was represented with arrays containing 101 images: the first twenty were gray (−0.2 s to 0.0 s), the next thirty were copies of the image presented to subjects in the experiment (0.0 to 0.3 s), and the remaining fifty-one (0.3 to 0.8 s) were gray. These were paired with output labels obtained by multiplying corresponding ERPs by one million to normalize their units (ERPs are typically in microvolts). The model was thereby optimized to produce outputs matching ERP waveforms in response to sequences of images shown in the experiment. Training was performed for 1000 epochs with a batch size of 8. The adaptive moment estimation optimizer was used with default parameters (0.001 learning rate, 0.9 beta-1, 0.99 beta-2) and mean-squared error loss. Input arrays, computational model, CNN vector outputs, and output labels are illustrated in Figure 2.

**Figure 2.**
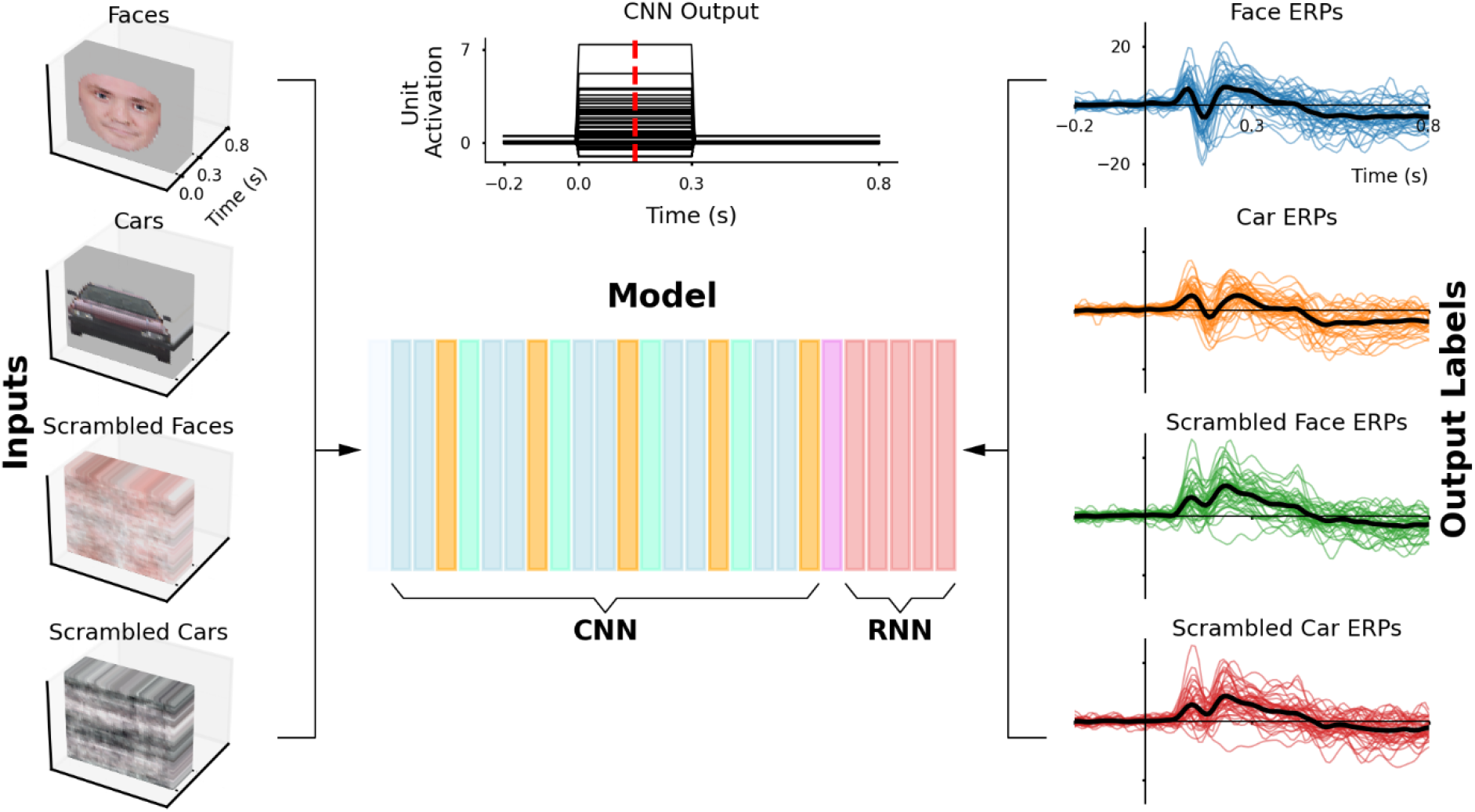
Supervised learning paradigm. Inputs on the left-hand side were three-dimensional representations of image time-sequences presented in the visual ERP experiment. Four stimulus categories were presented: faces, cars, scrambled faces and scrambled cars, each comprising 40 unique images. The computational model is illustrated conceptually in the center. It consisted of a three-dimensional convolutional neural network (CNN) connected to a recurrent neural network (RNN). The CNN transformed each image of the input time-sequence into a vector representation. An example of CNN outputs for one input image sequence is shown at the top, with a red dashed line transecting the vector representation during stimulus image presentation from 0.0 to 0.3 s. The model was trained end-to-end, with output labels provided from ERP waveforms recorded from a cohort of human subjects as they viewed the same stimuli depicted by the inputs. Grand-average ERPs are identified with thick black traces.

#### 2.2.3. Cross validation

The model architecture was evaluated with ten-fold cross validation. In each fold, four different pairs of input arrays and output labels from each stimulus category (F, C, SF and SF) were set aside to be used as validation data. A fresh model was instantiated and trained using the remaining 144 input-label pairs. After training, models were used to predict ERP waveforms in response to held-out validation data, producing a complete set of 160 cross-validation (CV) ERPs over ten folds. Overall performance was evaluated by comparing how well these model outputs correlated with real ERPs, displayed on Figure 3. After cross-validation, a final model was trained using all of the available data. The behavior of this final model was then analyzed to investigate how it generates outputs replicating ERP waveforms in response to sequences of visual input.

**Figure 3.**
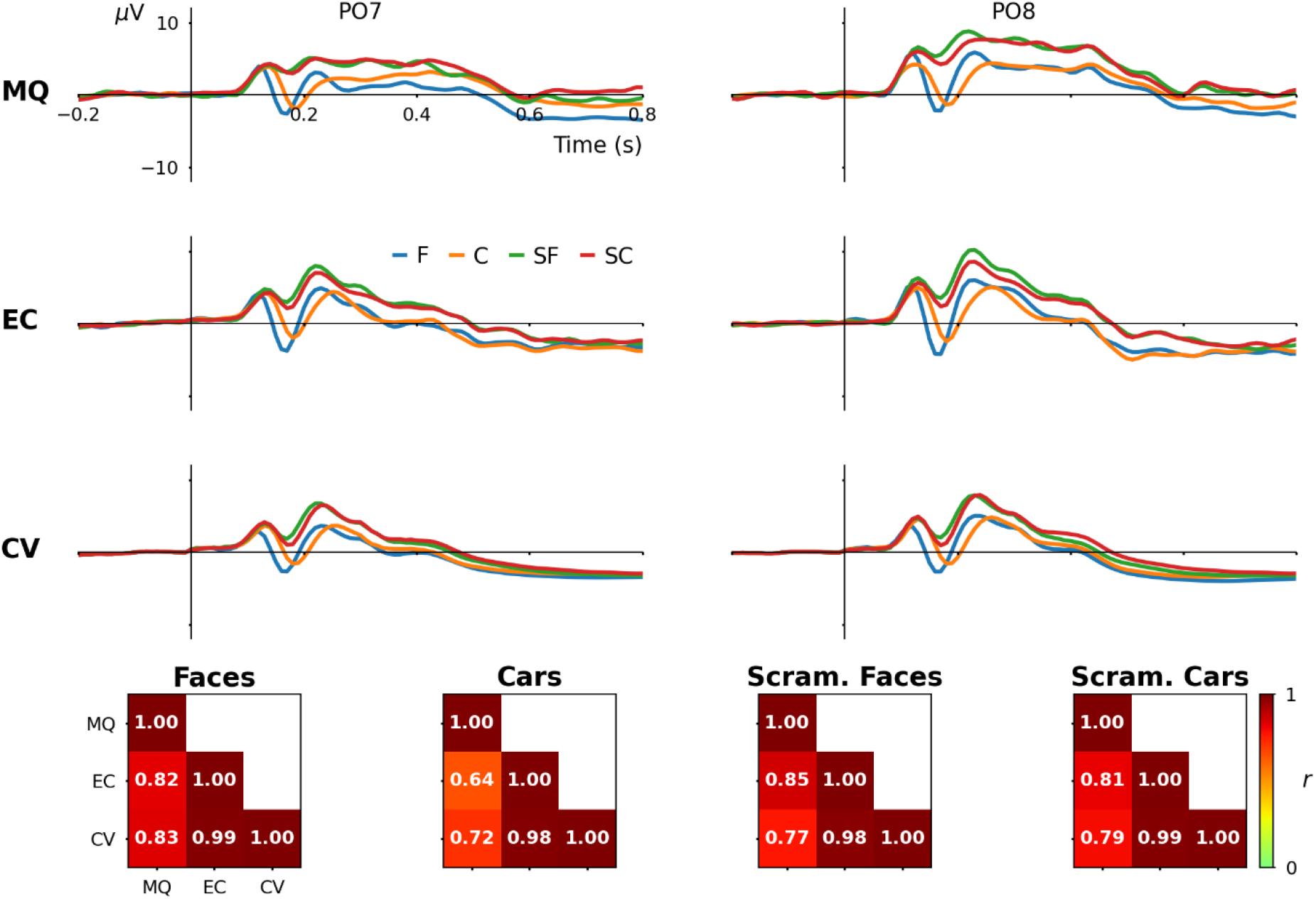
Grand-average event-related potential waveforms from two human cohorts and computational model. The first row displays data from the Macquarie University (MQ) validation study (n = 16); the second row shows data from the ERP CORE (EC) dataset (n = 40); the third row presents outputs from model cross-validation (CV) cases (n = 40). The bottom row depicts correlation (r) across the three datasets for face (F), car (C), scrambled face (SF) and scrambled car (SC) ERP waveforms. For all correlations, p < 0.001.

#### 2.2.4. Model analysis

##### 2.2.4.1. Principal component analysis of image vector representations

Principal component analysis was used to reduce the dimensionality of CNN vector representations from 240 to 2 for visualization. The vector representing each stimulus image was retrieved from a single time-slice of the CNN output, taken from a time-point when the stimulus image was shown, illustrated by the dashed line in the upper middle panel of Figure 2. These vector representations were generated by the final model that had been trained on the whole EC dataset. The singular value decomposition transform was determined from vector representations of EC stimuli. Both EC and synthetic image vector representations were then transformed into two principal components for visual comparison. These data are plotted in Figure 4.

**Figure 4.**
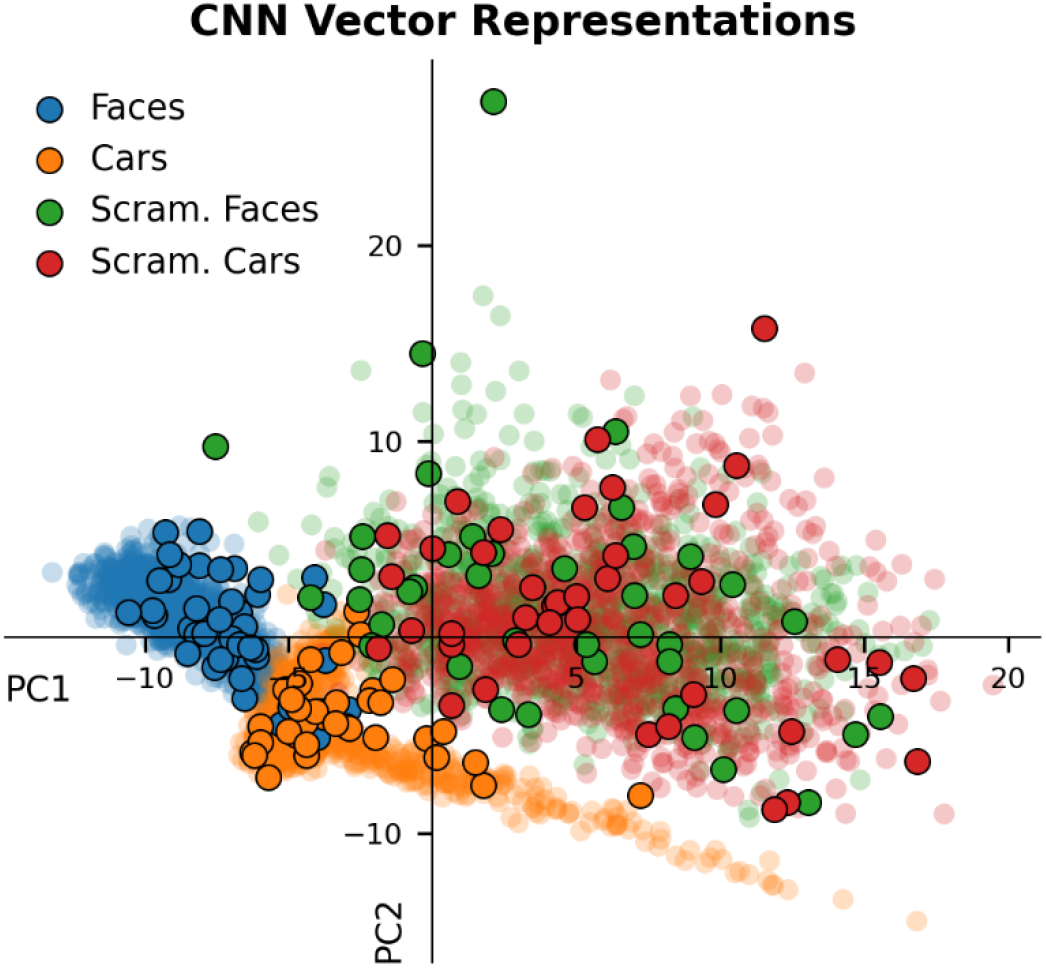
Principal component analysis of vector representations learned by the convolutional neural network. Each vector representation of 240 elements was transformed into two principal components for visualization. Real image data points (n = 40 for each image category) are shown by solid markers with black outlines. Markers representing synthetic image data points (n = 1000 for each category) are slightly transparent.

##### 2.2.4.2. Classification analysis of image vector representations

To further evaluate vector representations produced by the CNN section, a logistic regression classifier was trained on vector representations from 160 real EC images and then tested on those from 4000 synthetic images. This was treated as a one-versus-rest multi-class classification problem, with four classes of images represented by the vectors. Comparing classification performance across the four categories of images provides information about how robustly these are represented by the 240 element vectors. For example, classes with high accuracy occupy a designated region in multidimensional space and have distinctive vector representations, whereas classes with low accuracy overlap in vector space and have less distinctive vector representations. Within the described end-to-end training paradigm, robustness of these vector representations will influence how well model outputs match ERP labels.

##### 2.2.4.3. Representational similarity analysis of model behavior and ERPs

Representational similarity analysis (RSA) ^27^ was applied to examine final model behavior relative to EC and MQ ERP waveforms. This is shown in Figure 5. Activations were obtained from the model at multiple levels: CNN output, RNN layer 1 to 4, and model output. Representational dissimilarity matrices (RDMs) were computed across the full epoch and also across five-sample sliding windows with step size of one. Correlations among both sets of ERP data and model activation RDMs were calculated. The RSA approach allows us to compare how model activations and neurophysiological signals vary in response to the same set of stimuli.

**Figure 5.**
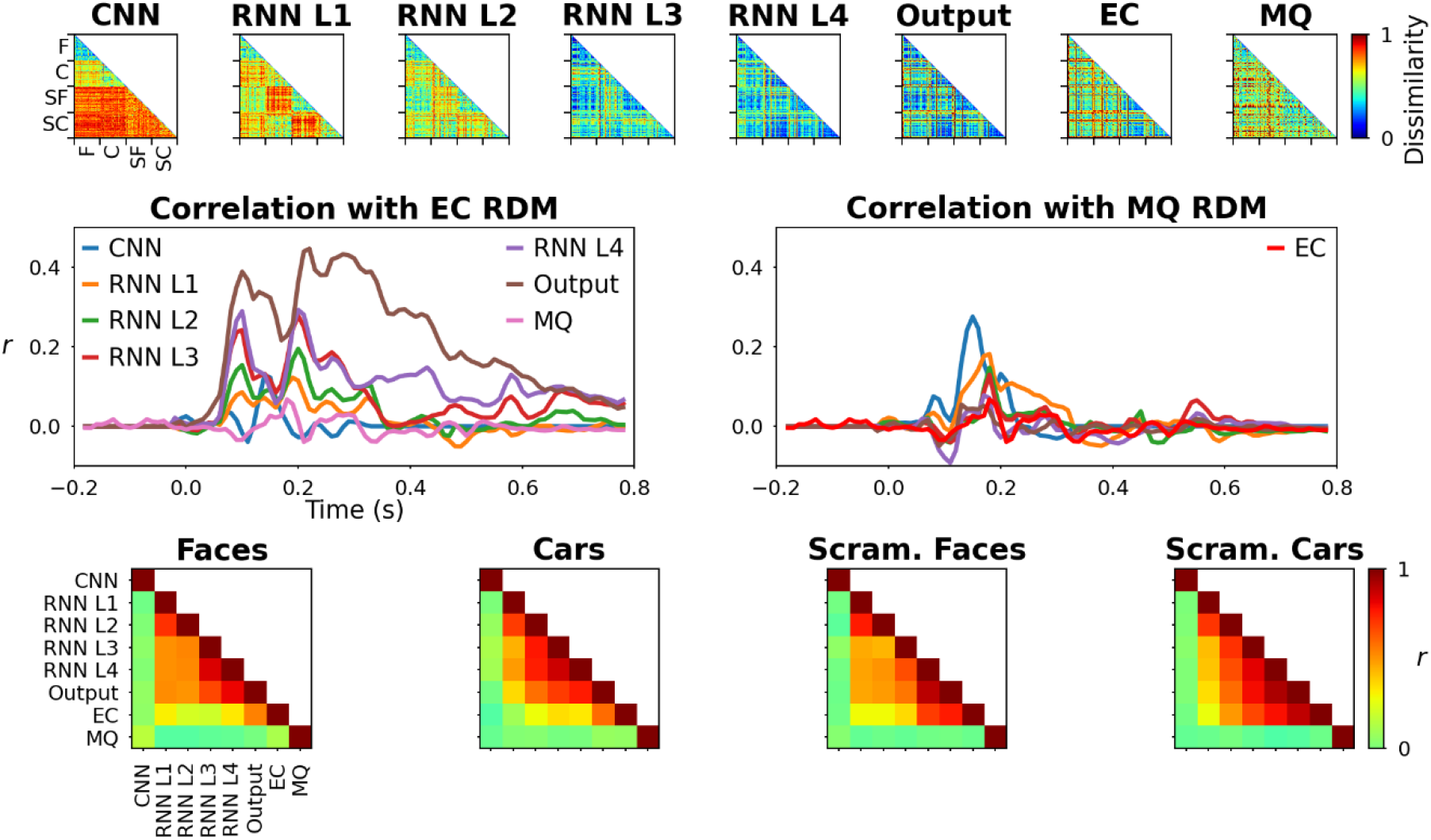
Representational similarity analysis. The top row shows representational dissimilarity matrices (RDMs) from CNN outputs, four RNN hidden layers (L1-L4), model output, ERP CORE (EC) ERPs and validation study (MQ) ERPs. These 160 by 160 matrices were computed by comparing 40 samples of four stimulus categories: faces (F), cars (C), and scrambled faces (SF) and cars (SC). The middle-left panel shows correlation between model and MQ with EC RDMs calculated from a five-sample sliding window with step size of one. The middle-right panel displays similar time-wise correlation between model and EC RDMs with the MQ RDMs. The bottom row displays correlation between RDMs in the top row by image category.

##### 2.2.4.4. Analysis of model RNN hidden units associated with face and car images

Hidden unit activations from the RNN section before the output were categorized as being associated mostly with face images, car images, or neither, according to the stimuli that elicited their maximal response. This was determined by integrating hidden unit activations over time and classifying them according to the stimulus category that maximized this. Three class separation was justified by the patterns of ERP waveforms and image vector representations that suggested overlap between the two types of scrambled images. Therefore, responses to both sets of scrambled images were grouped together and this analysis concentrated on differences between processing of face and car images. This data is plotted in Figure 6.

**Figure 6.**
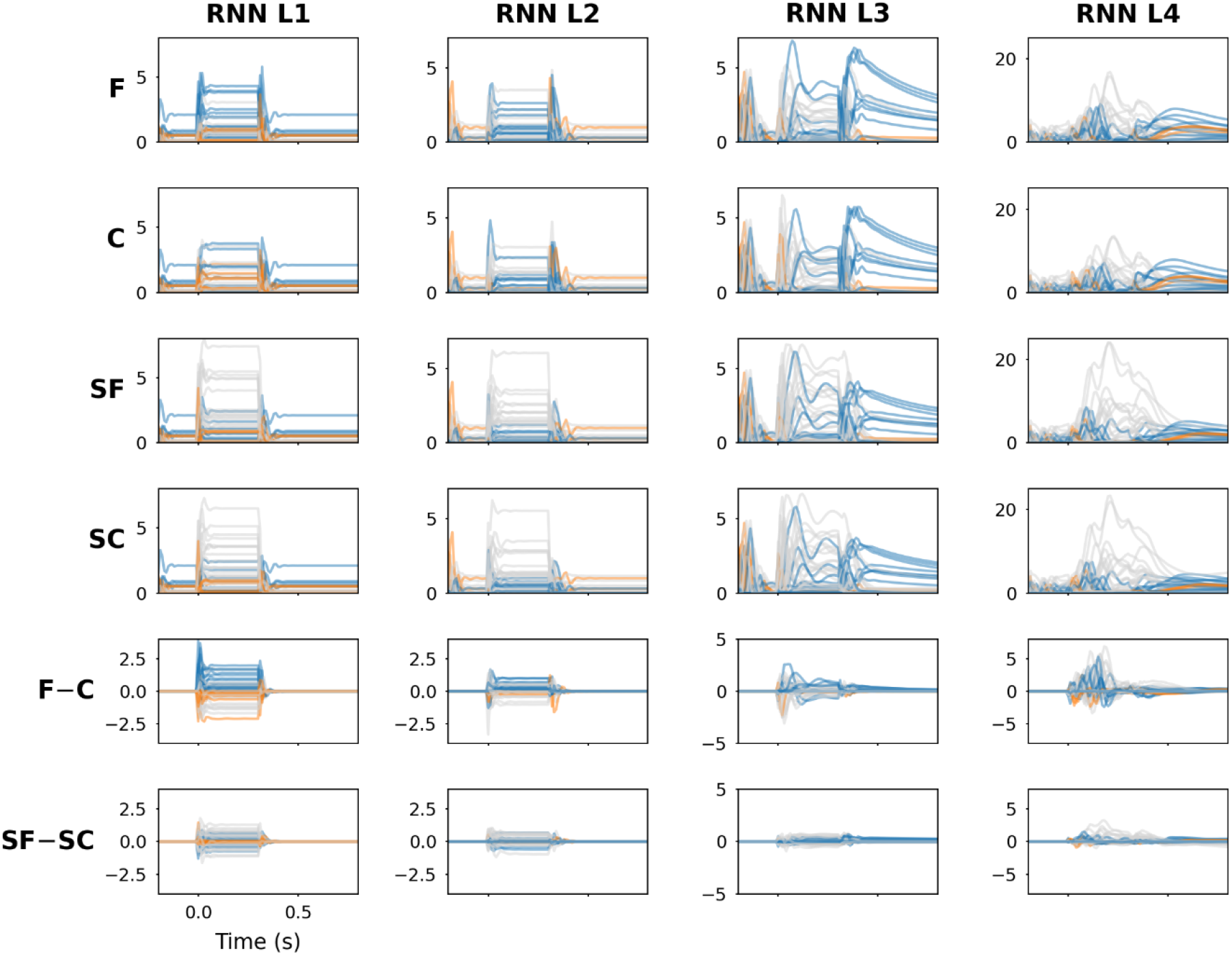
Hidden unit activations from the recurrent neural network section of the model. Units maximally sensitive to face stimuli are colored blue, those most sensitive to car stimuli are colored orange, and the rest are colored gray. Responses to faces (F), cars (C), scrambled faces (SF) and scrambled cars (SC) are shown, as well as waveform subtractions, F − C and SF – SC. Layer 4 activations were directly mapped onto model outputs representing ERP waveforms, so are considered to reflect candidate patterns of neural activity responsible for shaping ERP morpholohy.

#### 2.2.5. Simulated experiments

We ran simulated experiments using the final model. Synthetic images similar to those in the original ERP CORE experiment were used in these simulations. Artificial face and car images were generated with GANs and phase-scrambled to produce four image categories, each with 1000 samples. These were presented to the model in four conditions: upright, inverted, low-pass filtered (LF) and high-pass filtered (HF). Image filtering was performed in the frequency domain using a filter mask containing a circle with 5 px radius smoothed with a 5 by 5 px average filter. For the low-pass filter mask, the center circle, intersecting low spatial frequencies, was equal to 1, whereas outside the center circle, intersecting higher spatial frequencies, was equal to 0. The high-pass filter mask was an inverted copy of the low-pass filter mask; i.e., 0 at the center and 1 outside the center. Model outputs were inspected and difference waveforms face minus car (F – C) and scrambled face minus scrambled car (SF – SC) were computed. The results from these experiments are plotted in Figure 7. The MQ dataset was collected in a follow-up experiment to evaluate the results of simulations involving inverted images, the results of which are shown in Figure 8.

**Figure 7.**
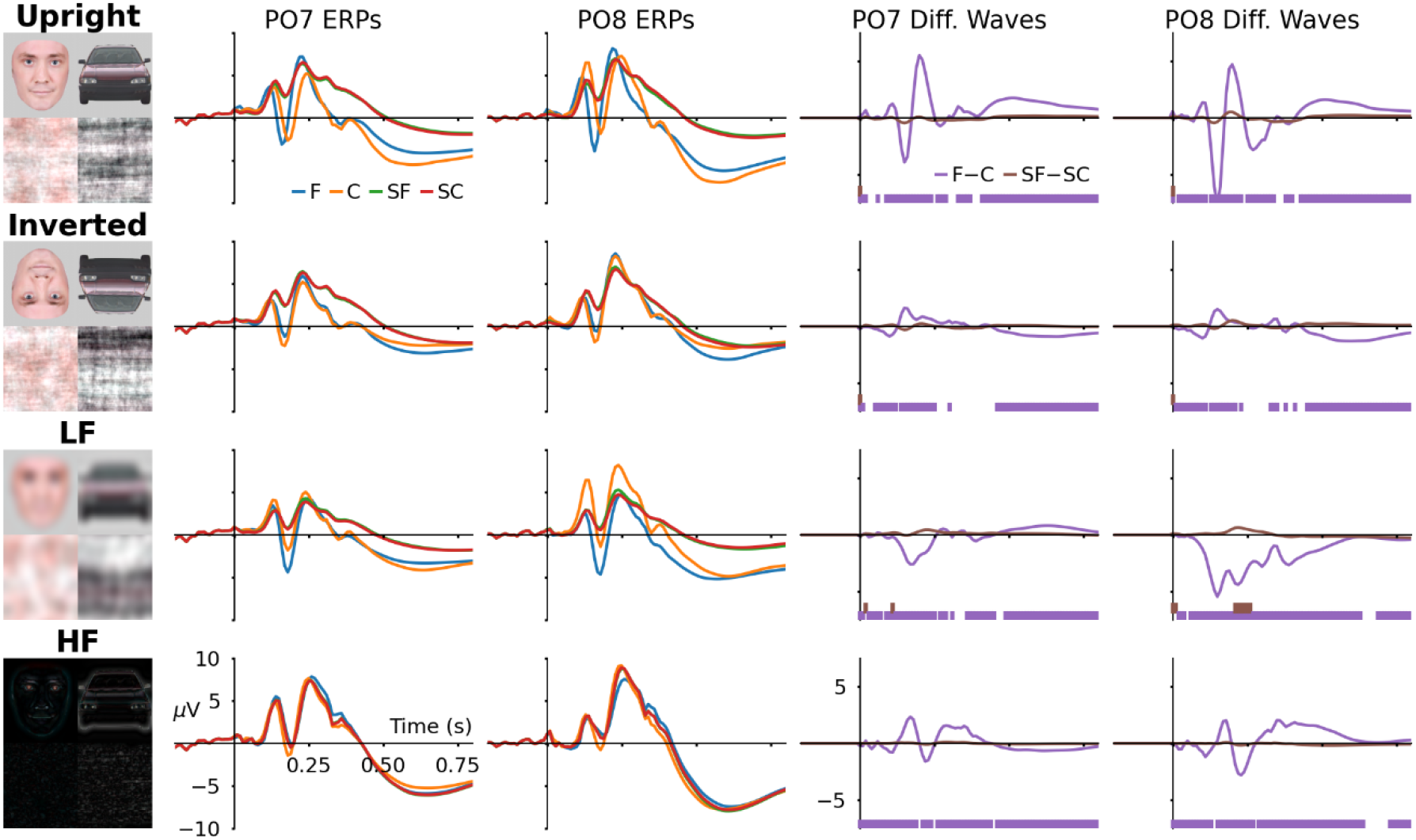
Simulated experiments performed with the model and synthetic images. One thousand artificial images of faces (F), cars (C), scrambled faces (SF) and scrambled cars (SC) were formatted as input arrays and fed into the model to predict corresponding ERP waveforms reflected by model outputs. Three manipulations were performed on these sets of images: they were inverted, low-pass filtered (LF) and high-pass filtered (HF). Time-points with statistically significant (p < 0.001, with Bonferroni corrections) differences between sets of model outputs used to compute difference waveforms are illustrated with colored bars below the difference waveform plots on the right.

**Figure 8.**
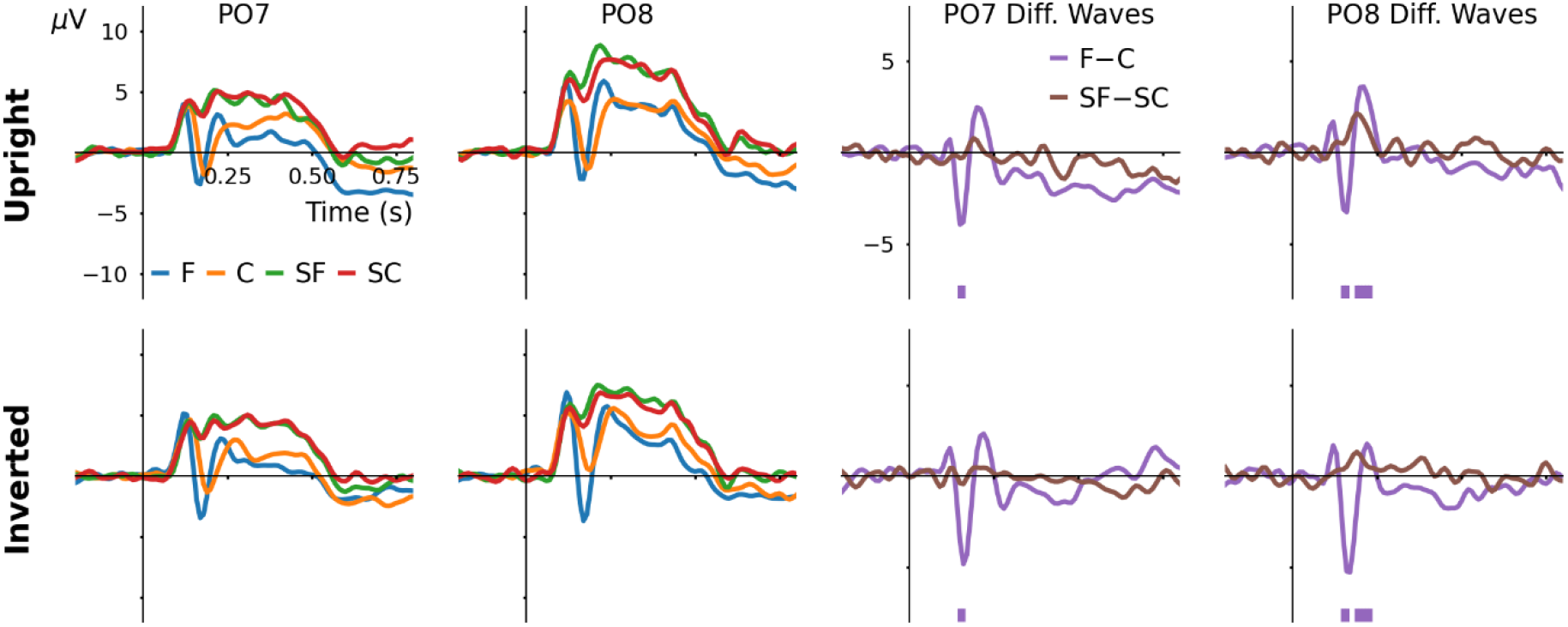
Validation experiment performed in a cohort of human subjects. The top row displays validation study (MQ) data from Figure 3. Upright and inverted images of faces (F), cars (C), scrambled faces (SF) and scrambled cars (SC) were presented to sixteen healthy adult subjects to evoke these ERP waveforms. Statistically significant results from timewise Mann-Whitney U-tests (p < 0.001, with Bonferroni corrections) comparing the two sets of ERPs used to produce difference waveforms are annotated below difference wave plots in their respective colors.

### 2.3. Statistical analysis

Pearson’s correlation coefficient (r) was used to compare ERPs from EC, MQ and CV datasets (Figure 3) and RDM matrices from CNN outputs, RNN layers and both sets of ERPs (Figure 4). Difference waveforms in Figure 7 and Figure 8 were evaluated by comparing the two associated sets of ERP waveforms with two-tailed Mann-Whitney U-tests at every time point, followed by Bonferroni corrections for multiple comparisons. The threshold for statistical significance was set conservatively for all statistical tests, with an alpha value of 0.001.

### 2.4. Software

Python 3 was used with Matplotlib 3.5.2, NeuroRA 1.1.6.8, NumPy 1.22.4, OpenCV-Python 4.6.0.66, Scikit-Learn 1.1.1, SciPy 1.8.1, and TensorFlow 2.9.1. The model and code will be shared in a public repository before publication.

## 3. Results

### 3.1.1. Model outputs are strongly correlated with real ERP waveforms

Grand-average ERPs from validation study (MQ), ERP CORE (EC) and model cross-validation output (CV) datasets are plotted in the upper three rows of Figure 3. Qualitatively, these three sets of ERPs are in close agreement. Correlation coefficients relating each pair of waveforms are provided in the bottom row of Figure 3, separated by stimulus category. Across all stimulus responses, CV and EC waveforms were highly correlated (r = 0.98, p < 0.001), CV and MQ were slightly less correlated (r = 0.78, p < 0.001) similar to EC and MQ waveforms (r = 81, p < 0.001). Individual ERPs from each dataset are plotted in supplementary figures S1-S4.

### 3.1.2. Learned image vector representations are specific to known objects

CNN vector representations transformed into two principal components are visualized in Figure 4. Faces and cars occupy distinct regions in this space whereas their scrambled counterparts overlap. A logistic regression classifier trained on full-size (240 element) vector representations of real images and tested on vectors representing synthetic images achieved an overall training accuracy score of 1.0 and testing accuracy score of 0. 802. Evaluating test set accuracy by image category: faces 1.0, cars 0.995, scrambled faces 0.523 and scrambled cars 0.689. In terms of misclassifications, four car image vectors were classified as scrambled faces and one was classified as scrambled car (see supplementary figure S5 for details). Scrambled face image vectors were incorrectly classified as face 7, car 27, and scrambled car 443 times. Vectors representing scrambled cars were misclassified as face 3, car 61, and scrambled face 247 times.

### 3.1.3. Representational similarity with neural data increases along model hierarchy

Model behavior was compared with EC and MQ responses using representational similarity analysis (RSA) ^27^. These analyses are visualized in Figure 5. Comparing the lower triangle of representational dissimilarity matrix (RDM) data displayed in the top row of Figure 5 with those of the EC dataset produced correlation coefficients of −0.017 (p = 0.0571) for CNN, 0.183 for RNN layer 1, 0.293 for RNN layer 2, 0.367 for RNN layer 3, 0.485 for RNN layer 4, 0.668 for model output; where not stated, p < 0.001. Comparing model activations with MQ data produced correlation coefficients of 0.018 (p = 0.04) for CNN, 0.014 (p = 0.11) for RNN layer 1, 0.029 (p = 0.001) for RNN layer 2, 0.012 (p = 0.18) for RNN layer 3, −0.013 (p = 0.13) for RNN layer 4, 0.028 (p = 0.0017) for model output. The two RDMs computed from real neural data (i.e., EC vs. MQ) had r = 0.021 (p = 0.0172).

When compared over time, patterns of correlation between model RDMs from the RNN section and EC RDMs tend towards displaying twin peaks at approximately 90 ms and 200 ms post stimulus onset. In contrast, CNN outputs displayed a single peak correlation at 140 ms (r = 0.128, p < 0.001). For both overall RDMs and time-wise RDMs these correlations were proportional to layer proximity to the model output; i.e., the RDM from model output was most strongly correlated, whereas the RDM from CNN outputs were least strongly correlated with the EC RDM data. Compared with the MQ RDM data across time, CNN activity peaked in correlation at 150 ms (r = 0.275, p < 0.001), RNN layers 1-3 peaked at 180 ms (r = 0.182, r = 0.147, r = 0.13; all p < 0.001), RNN layer 4 peaked in correlation at 170 ms (r = 0.0781, p < 0.001), and model output and MQ RDMs correlated with a peak at 0.13 s (r = 0.0545, p < 0.001). Correlation between MQ and EC RDM data peaked at 180 ms (r = 0.0676, p < 0.001).

### 3.1.4. Recurrent units show differences between face and car image processing

Hidden unit activations from the RNN section are shown in Figure 6. Responses to face, car, scrambled face and scrambled car stimuli are displayed in the top four rows. Face minus car and scrambled-face minus scrambled-car responses are plotted in the bottom two rows. Traces are colored according to which stimulus category elicited maximum responses from each hidden unit: face (blue), car (orange) or scrambled (gray). Substantial differences between responses to face and car stimuli are observed from outputs of layer one, the first transformation of CNN vector outputs. Differences between responses to face and car images are apparent as information propagates through the network to layer four. In comparison, differences between scrambled face and scrambled car inputs are less pronounced at every layer. Activations from layer four units are combined in a linear superposition to produce model outputs. These are therefore conceptually similar to neural sources that contribute to scalp-recorded ERP signals. The same layer four units are activated by both face and car stimuli, however, the magnitude of responses to faces tended to be greater than those of cars.

### 3.1.5. Simulations suggest patterns of differences between inverted, low-frequency and high-frequency stimuli

Results from four simulated experiments are shown in Figure 7. Upright images produced an N170 response to faces that was earlier and greater amplitude than that of car images. Difference waveforms between upright face and car responses highlight this feature and its laterality towards the right hemisphere (channel PO8). Upright scrambled images did not produce significant differences in model outputs. Inverting car and face images inverted the pattern of ERP differences between their model responses. Low-frequency face stimuli elicited more negative amplitude model output relative to car stimuli, whereas high-frequency produced phasic differences in model output deflections. Inverted, low-frequency and high-frequency scrambled images did not produce substantially different outputs from the model.

### 3.1.6. Validation study is partly consistent with model simulations

Event-related potentials obtained from the MQ validation experiment with inverted images are plotted in Figure 8. The face minus car ERP difference waveform produced by responses to upright images has more well-defined alternating polarity morphology, whereas that produced by inverted images shows a relatively pronounced negative peak and diminished positive peak. These differences reflect changes to ERP waveforms evoked by face and car images when they are inverted. In contrast, difference waveforms produced by subtracting responses to scrambled car images from those of scrambled face images do not change notably when the images are upside down. Model outputs in response to inverted scrambled images are consistent with these observations, whereas model outputs in response to inverted face and car images are not.

## Discussion

The aim of developing this CNN-RNN model was to explore whether it can reproduce ERP responses evoked by images of faces, cars, scrambled faces and scrambled cars. This was demonstrated with cross-validation model outputs shown in Figure 1 and supplementary figures S1-S4. Maximum correlation was observed between model outputs and ERP CORE data, which is unsurprising given that samples from this dataset were used to develop the models. The validation study dataset correlated less highly with cross-validation model outputs. This reduction is considered to reflect data variability in ERP experiments conducted in separate labs, with different participants, using slightly different equipment. However, the still relatively high correlations should be taken as an indication of the similarity between the datasets.

Vector representations learned by the CNN section distinguished between face and car stimuli but not their phase-scrambled equivalents. This is seen in two-dimensional principal component space plotted in Figure 4, and also from the results of vector classification. More accurate classification of face and car image vectors compared with scrambled image vectors suggests that the model only needed to identify these three categories to reliably generate output waveforms matching ERPs. This can be explained by the fact that ERP waveforms elicited by scrambled face and scrambled car images were effectively indistinguishable. This indicates that semantic content within images is more consequential than color for producing model outputs and ERP waveforms, given that the scrambled images contained the same colors as unscrambled images in unrecognizable configurations. The research literature on N170 sensitivity to faces, other objects and non-objects is consistent with these observations ^4,6,9,28^.

Representational similarity analysis ^27^ was used to compare neural data from ERP CORE (EC) and the validation study (MQ) with final CNN-RNN model activations in Figure 5. Representational dissimilarity matrices (RDMs) were computed to quantify correlation between model activations at multiple levels in response to 160 different input stimuli across the four categories. When calculated over the whole epoch, correlation between model and EC RDMs increased with proximity to the model output. In contrast, model and MQ RDMs showed negligible correlation when evaluated over the whole epoch. However, when evaluated timewise over five-sample sliding windows, significant correlations between model activations and both MQ and EC RDMs emerged, principally between 150 and 180 ms, suggesting that meaningful variance in the neural responses to stimulus images is concentrated within this time period. Correlations between EC and model activations tended to be higher than some previous reports ^29,30^, presumably due to the supervised learning paradigm applied to constrain our model to replicate these ERP waveforms. Although correlations between model activations and MQ data, and EC and MQ data were more closely aligned with previous results of RSA ^29,30^.

Hidden units in the RNN section of the model demonstrated different patterns of activity in response to face and car stimuli, illustrated in Figure 6. At the fourth hidden layer, the majority of these differences occurred before 200 ms after stimulus onset. Activations of these units are weighted together to generate model outputs and are therefore considered analogous to the activity of neural sources. We can see that the same units were activated by all stimuli, although the magnitudes of those activations were dependent upon stimulus category, with face images tending to produce larger amplitude responses. Interpreted within the context of the debate regarding face versus familiar-object specificity of N170, these findings support the view that the underlying source activity is sensitive to familiar objects rather than being specific to a particular category of objects. This should not be overstated, however, as there is inherent difficulty in separating neighboring, temporally-overlapping sources from ERP signals. Furthermore, this may reflect a tendency for the RNN’s hidden units to not be stimulus-specific per-se (i.e., not solely responding to a specific category of stimuli), but instead have their amplitudes modulated by different types of stimuli.

Synthetic images generated by GANs were realistic-looking and elicited reasonable responses from the model. These were created to evaluate model performance on a large set of unseen images from the same distribution as the real images used in EC and MQ experiments. Synthetic face images produced an N170 peak with greater amplitude at channel PO8 relative to PO7. Vector representations produced by the model in response to synthetic images (semi-transparent markers in Figure 4) were also clustered according to their intended category. From the simulation results illustrated in Figure 7, differences between face and car images change with each type of image manipulation, whereas differences between scrambled images remain constant. This suggests that when face and car images are sufficiently distorted the model ceases to output waveforms comparable with those in the training data. It is not unreasonable to suspect that changes to the appearance of images might similarly influence ERP waveforms ^31–33^.

Results from the validation experiment in Figure 8 do not concur with model responses to inverted face and car images (second row of Figure 7). The face N170 peak seen in real ERPs is enlarged and slightly delayed when images are inverted ^5,7,8^, whereas model outputs to inverted faces were attenuated. Comparing MQ ERPs and model outputs in response to inverted face and inverted car images (supplementary figure S6) highlights these differences. In contrast, validation experiment ERPs in response to scrambled images were relatively unaffected by inversion, which is consistent with model outputs. This is unsurprising because random phase-scrambled images do not have a “right way up”, therefore the model (and presumably the human brain) makes no distinction whether they are upright and or inverted. Considering the variable success of these predictions, we cannot assume that model outputs in response to low- and high-pass filtered images are accurate representations of ERPs that would be elicited by these images in real experiments. Nevertheless, it is likely that these image manipulations would cause some changes to ERP morphology ^5,7,8^.

In future work this model could be fine-tuned with data acquired from experiments investigating more granular aspects of human face processing such as ambiguity ^34^, emotion ^35^, familiarity ^36,37^, racial congruence ^38^, or wearing of partial facial coverings ^39,40^ to evaluate how these affect model behavior. Doing so would provide additional analyses to assist in interpreting computational principles underlying sensitivity of brain responses to different manipulations of face stimuli. Furthermore, this modeling approach could be applied in other visually-evoked potential contexts as a model of ERP generation.

## Conclusion

The CNN-RNN model reliably captured some of the key features of ERP responses to face, car and phase-scrambled images. These include N170 sensitivity to face images lateralized to the right hemisphere. Different latent states observed from the model became increasingly correlated with ERP responses as they were transformed from vector representations through to model output signals. This is somewhat analogous to visual information processing from the retina through to higher-order occipital cortex. However, model predictions in response to inverted face images were inconsistent with expectations and our validation experiment, highlighting an important limitation. The model could not reliably reproduce unseen patterns of neural activity in response to image manipulations. Nevertheless, further development and application of this model may be beneficial for analyzing subtler aspects of face perception and generally to explore the computational principles underlying visually evoked potentials.

## Author contributions

JAO: Conceptualization, Methodology, Software, Formal analysis, Data curation, Writing - original draft, Visualization, Supervision, Funding acquisition

JW: Conceptualization, Methodology, Software, Formal analysis, Data curation, Writing - review & editing, Supervision

AC: Validation, Investigation

JB: Validation, Investigation

TH: Validation, Investigation

FA: Methodology, Software, Formal analysis

PFS: Resources, Writing - review & editing, Supervision, Project Administration, Funding acquisition

## Acknowledgements

This work was supported in part by an International Brain Research Organization (IBRO) Exchange Fellowship awarded to JAO to train with PFS. The computer and graphics processing unit used to train the models were purchased with a grant from the Research Institute of Rangsit University (grant number 90/2561).

## Data availability

The event-related potential recordings and image stimuli used to develop the model are available from ERP CORE which can be accessed from https://osf.io/thsqg/.

## Ethical approval

The validation study conducted at Macquarie University was reviewed and approved by the Human Research Ethics Committee.

## References

1. Leopold, D. A. & Rhodes, G. A Comparative view of face perception. J. Comp. Psychol. 124, 233–251 (2010).

2. Zagury-Orly, I., Kroeck, M. R., Soussand, L. & Cohen, A. L. Face-Processing Performance is an Independent Predictor of Social Affect as Measured by the Autism Diagnostic Observation Schedule Across Large-Scale Datasets. J. Autism Dev. Disord. 52, 674–688 (2022).

3. Behrmann, M. & Avidan, G. Congenital prosopagnosia: face-blind from birth. Trends Cogn. Sci. 9, 180–187 (2005).

4. Osborne, K. J., Kraus, B., Curran, T., Earls, H. & Mittal, V. A. An Event-Related Potential Investigation of Early Visual Processing Deficits During Face Perception in Youth at Clinical High Risk for Psychosis. Schizophr. Bull. 48, 90–99 (2022).

5. Wehrman, J., Sörensen, S., De Lissa, P. & Badcock, N. A. EPOC outside the shield: comparing the performance of a consumer-grade EEG device in shielded and unshielded environments. Biomed. Phys. Eng. Express 7, 025010 (2021).

6. Rossion, B. & Caharel, S. ERP evidence for the speed of face categorization in the human brain: Disentangling the contribution of low-level visual cues from face perception. Vision Res. 51, 1297–1311 (2011).

7. Bossi, F. et al. Theta- and Gamma-Band Activity Discriminates Face, Body and Object Perception. Front. Hum. Neurosci. 14, 74 (2020).

8. Bentin, S., Allison, T., Puce, A., Perez, E. & McCarthy, G. Electrophysiological Studies of Face Perception in Humans. J. Cogn. Neurosci. 8, 551–565 (1996).

9. Torriero, S. et al. FEF excitability in attentional bias: A TMS-EEG study. Front. Behav. Neurosci. 12, 333 (2019).

10. Ricciardelli, P., Ro, T. & Driver, J. A left visual field advantage in perception of gaze direction. Neuropsychologia 40, 769–777 (2002).

11. Richards, B. A. et al. A deep learning framework for neuroscience. Nat. Neurosci. 2019 2211 22, 1761–1770 (2019).

12. Richards, B., Tsao, D. & Zador, A. The application of artificial intelligence to biology and neuroscience. Cell 185, 2640–2643 (2022).

13. Lindsay, G. W. Convolutional Neural Networks as a Model of the Visual System: Past, Present, and Future. J. Cogn. Neurosci. 33, 2017–2031 (2021).

14. O’Reilly, J. A. Recurrent Neural Network Model of Human Event-related Potentials in Response to Intensity Oddball Stimulation. Neuroscience 504, 63–74 (2022).

15. O’Reilly, J. A., Angsuwatanakul, T. & Wehrman, J. Decoding violated sensory expectations from the auditory cortex of anaesthetised mice: Hierarchical recurrent neural network depicts separate ‘danger’ and ‘safety’ units. Eur. J. Neurosci. (2022) doi:10.1111/ejn.15736.

16. O’Reilly, J. A., Wehrman, J. & Sowman, P. F. A Guided Tutorial on Modelling Human Event-Related Potentials with Recurrent Neural Networks. Sensors 22, 9243 (2022).

17. O’Reilly, J. A. Modelling mouse auditory response dynamics along a continuum of consciousness using a deep recurrent neural network. J. Neural Eng. 19, (2022).

18. Shi, J., Wen, H., Zhang, Y., Han, K. & Liu, Z. Deep recurrent neural network reveals a hierarchy of process memory during dynamic natural vision. Hum. Brain Mapp. 39, 2269 (2018).

19. Xu, G., Ren, T., Chen, Y. & Che, W. A One-Dimensional CNN-LSTM Model for Epileptic Seizure Recognition Using EEG Signal Analysis. Front. Neurosci. 14, 1253 (2020).

20. Kappenman, E. S., Farrens, J. L., Zhang, W., Stewart, A. X. & Luck, S. J. ERP CORE: An open resource for human event-related potential research. Neuroimage 225, 117465 (2021).

21. Goodfellow, I. et al. Generative adversarial nets. Adv. Neural Inf. Process. Syst. 27, (2014).

22. Karras, T. et al. Training Generative Adversarial Networks with Limited Data. Adv. Neural Inf. Process. Syst. 2020-December, (2020).

23. Karras, T. et al. Analyzing and Improving the Image Quality of StyleGAN. Proc. IEEE Comput. Soc. Conf. Comput. Vis. Pattern Recognit. 8107–8116 (2019) doi:10.48550/arxiv.1912.04958.

24. Heusel, M., Ramsauer, H., Unterthiner, T., Nessler, B. & Hochreiter, S. GANs Trained by a Two Time-Scale Update Rule Converge to a Local Nash Equilibrium. Adv. Neural Inf. Process. Syst. 2017-Decem, 6627–6638 (2017).

25. Ioffe, S. & Szegedy, C. Batch normalization: Accelerating deep network training by reducing internal covariate shift. in 32nd International Conference on Machine Learning, ICML 2015 vol. 1 448–456 (International Machine Learning Society (IMLS), 2015).

26. Melinsca, M., Prentasic, P. & Loncaric, S. Retinal vessel segmentation using deep neural networks. in VISAPP 2015 - 10th International Conference on Computer Vision Theory and Applications; VISIGRAPP, Proceedings vol. 1 577–582 (2015).

27. Kriegeskorte, N., Mur, M. & Bandettini, P. Representational similarity analysis - connecting the branches of systems neuroscience. Front. Syst. Neurosci. 2, 4 (2008).

28. Rousselet, G. A., Pernet, C. R., Bennett, P. J. & Sekuler, A. B. Parametric study of EEG sensitivity to phase noise during face processing. BMC Neurosci. 9, 1–22 (2008).

29. Kiat, J. E., Hayes, T. R., Henderson, J. M. & Luck, S. J. Rapid Extraction of the Spatial Distribution of Physical Saliency and Semantic Informativeness from Natural Scenes in the Human Brain. J. Neurosci. 42, 97–108 (2022).

30. He, T., Boudewyn, M. A., Kiat, J. E., Sagae, K. & Luck, S. J. Neural correlates of word representation vectors in natural language processing models: Evidence from representational similarity analysis of event-related brain potentials. Psychophysiology 59, e13976 (2022).

31. Male, A. G. et al. The quest for the genuine visual mismatch negativity (vMMN): Event-related potential indications of deviance detection for low-level visual features. Psychophysiology 57, (2020).

32. Johannes, S., Münte, T. F., Heinze, H. J. & Mangun, G. R. Luminance and spatial attention effects on early visual processing. Cogn. Brain Res. 2, 189–205 (1995).

33. Lacroix, A. et al. The Predictive Role of Low Spatial Frequencies in Automatic Face Processing: A Visual Mismatch Negativity Investigation. Front. Hum. Neurosci. 0, 101 (2022).

34. Abubshait, A., Momen, A. & Wiese, E. Pre-exposure to Ambiguous Faces Modulates Top-Down Control of Attentional Orienting to Counterpredictive Gaze Cues. Front. Psychol. 11, 2234 (2020).

35. Petrucci, M. & Pecchinenda, A. The role of cognitive control mechanisms in selective attention towards emotional stimuli. Cogn. Emot. 31, 1480–1492 (2017).

36. Chauhan, V., Kotlewska, I., Tang, S. & Gobbini, M. I. How familiarity warps representation in the face space. J. Vis. 20, 18–18 (2020).

37. Wiese, H. et al. Familiarity Is Familiarity Is Familiarity: Event-Related Brain Potentials Reveal Qualitatively Similar Representations of Personally Familiar and Famous Faces. J. Exp. Psychol. Learn. Mem. Cogn. 48, 1144–1164 (2022).

38. Sessa, P. & Dalmaso, M. Race perception and gaze direction differently impair visual working memory for faces: An event-related potential study. Soc. Neurosci. 11, 97–107 (2016).

39. Noyes, E., Davis, J. P., Petrov, N., Gray, K. L. H. & Ritchie, K. L. The effect of face masks and sunglasses on identity and expression recognition with super-recognizers and typical observers. R. Soc. Open Sci. 8, (2021).

40. Calbi, M. et al. The consequences of COVID-19 on social interactions: an online study on face covering. Sci. Reports 2021 111 11, 1–10 (2021).

