## Supplemental Figures for "Neural correlates of face perception modeled with a convolutional recurrent neural network"

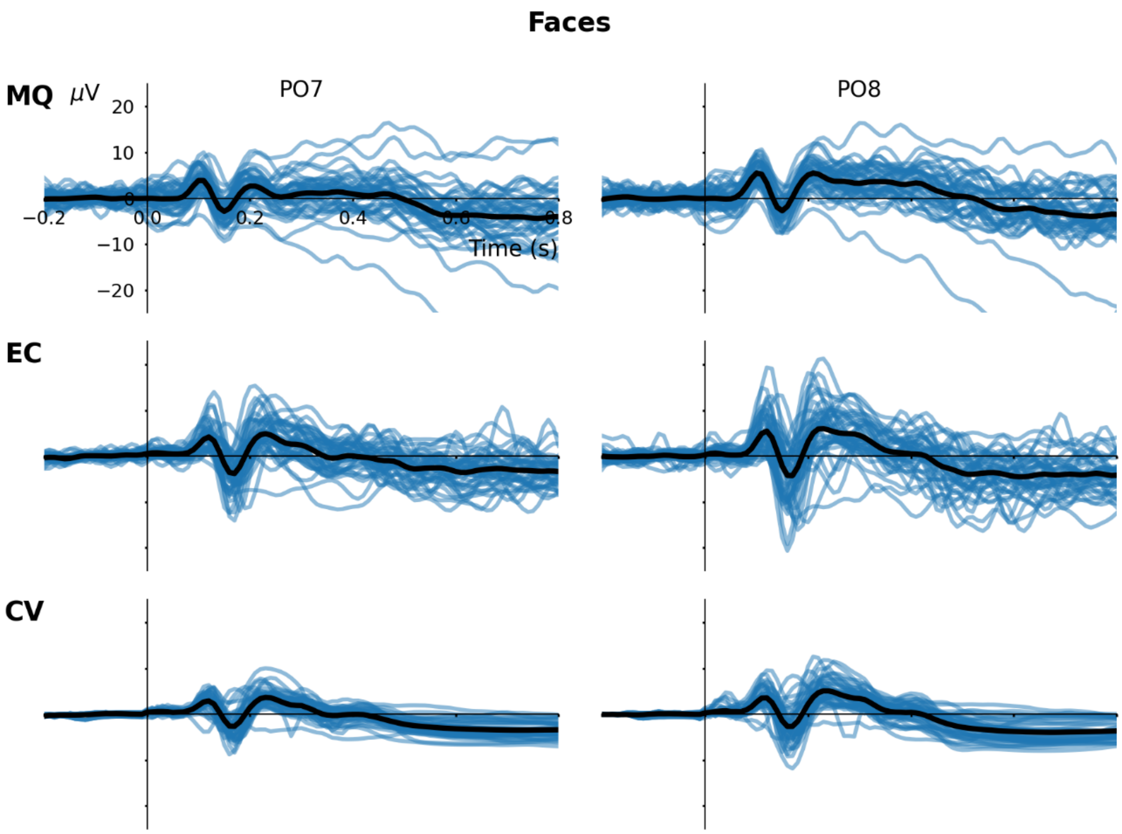


**Figure S1** Macquarie study grand-average (MQ), ERP CORE grand-average (EC) and model cross-validation (CV) waveforms from 40 unique face images.

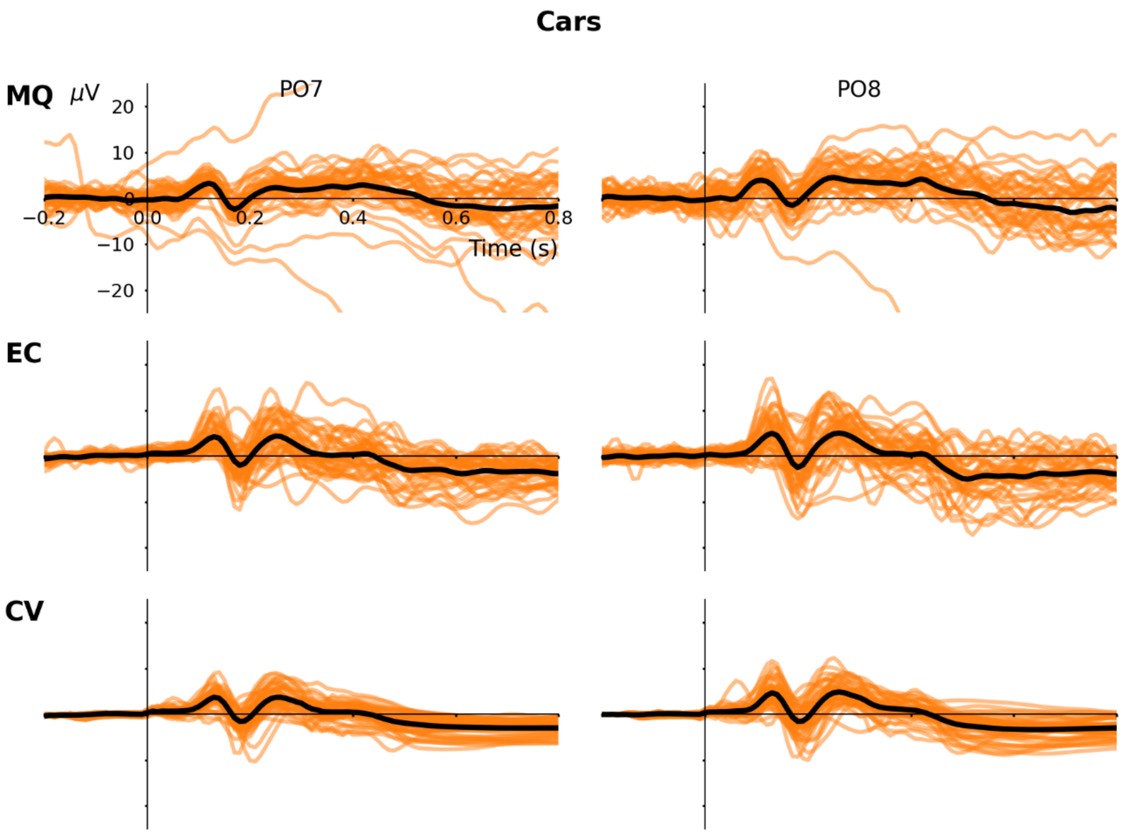


**Figure S2** Macquarie study grand-average (MQ), ERP CORE grand-average (EC) and model cross-validation (CV) waveforms from 40 unique car images.

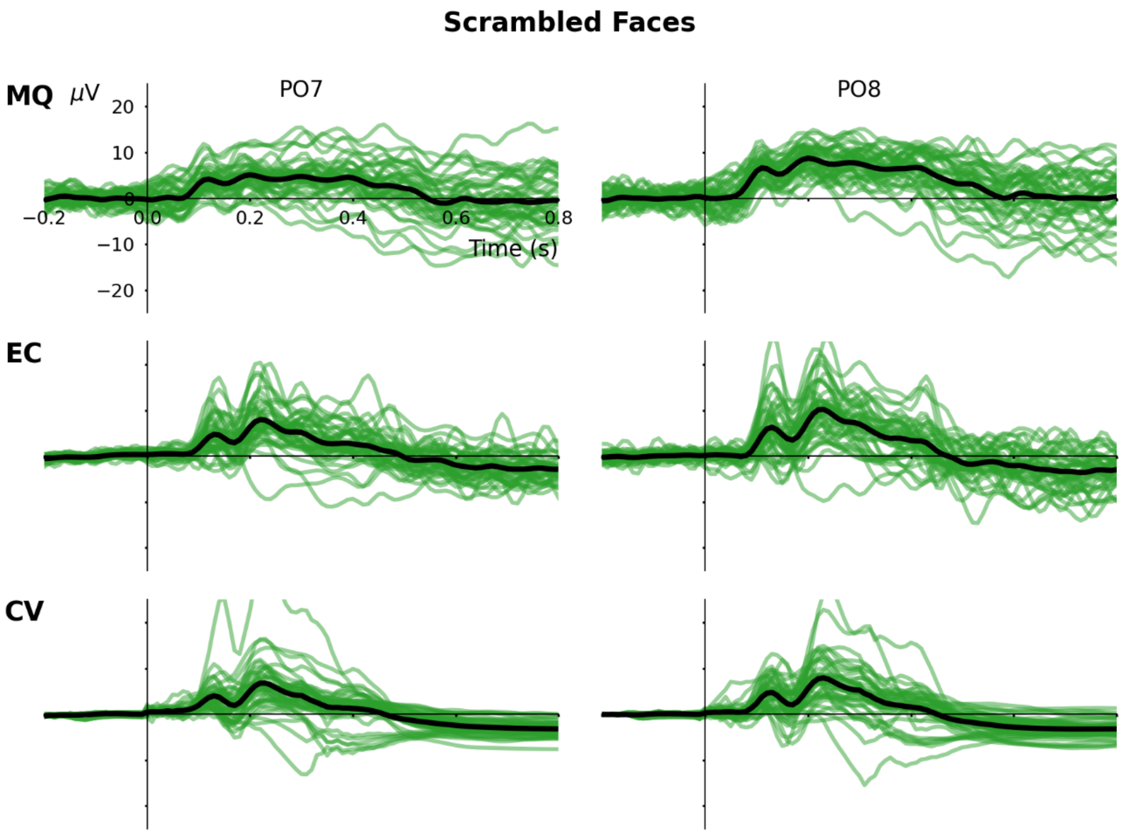


**Figure S3** Macquarie study grand-average (MQ), ERP CORE grand-average (EC) and model cross-validation (CV) waveforms from 40 unique scrambled face images.

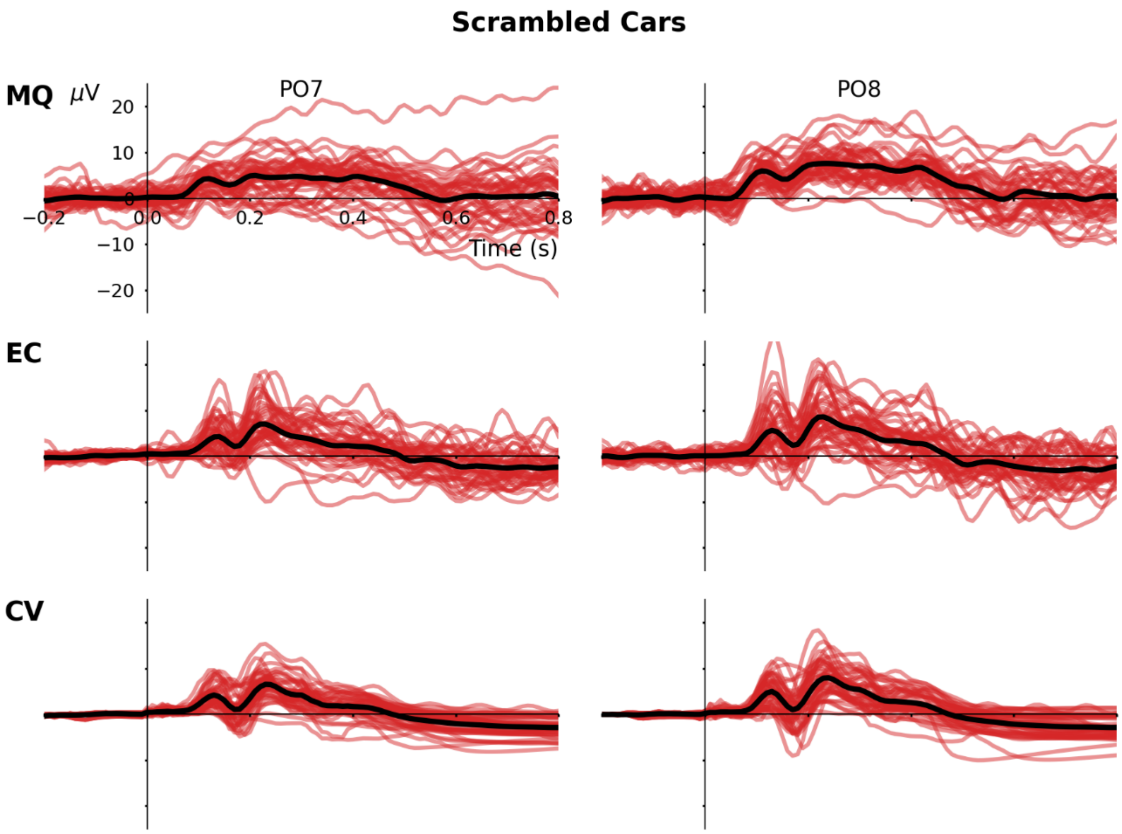


**Figure S4** Macquarie study grand-average (MQ), ERP CORE grand-average (EC) and model cross-validation (CV) waveforms from 40 unique scrambled car images.

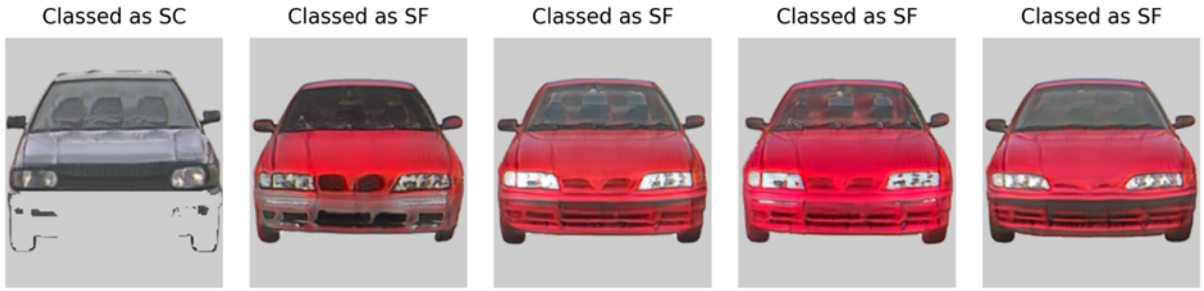


**Figure S5** Synthetic car images whose CNN vector representations were misclassified. The one on the left, a poorly constructed synthetic image, was classified as scrambled car (SC). The other three images were classified as scrambled face (SF) images.

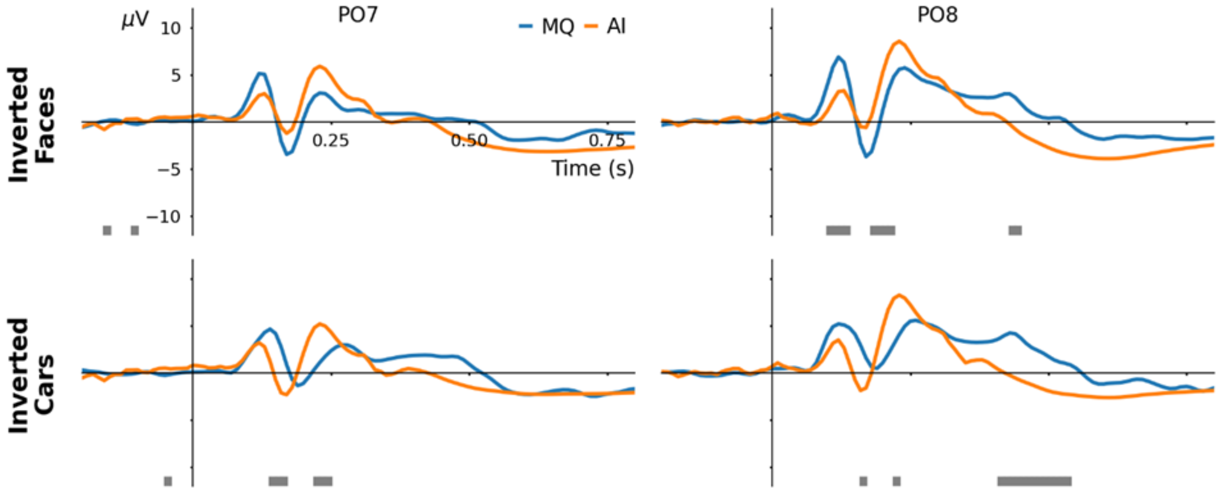


**Figure S6** Comparison between Macquarie validation study (MQ) and model output (AI) waveforms evoked by inverted face and car images. Below each panel, statistically significant time-points determined by bootstrapped Mann-Whitney U-tests are reported with grey bars ($p < 0.001$, with Bonferroni corrections). These tests were performed by randomly sampling model outputs in response to inverted synthetic images to compare with the equal sized sample of MQ ERPs in response to inverted real images ($n = 40$); 1000 permutations were performed and the average p-value at each time-point was calculated before applying Bonferroni corrections.
